# Impact of Group Size and Habitat Disturbance on Parasitic Infection in Free-ranging Proboscis Monkeys

**DOI:** 10.1101/2025.05.21.655443

**Authors:** Muhammad Nur Fitri-Suhaimi, Liesbeth Frias, Elke Zimmermann, Primus Lambut, Joseph Tangah, Henry Bernard, Vijay Subbiah Kumar, Ikki Matsuda

## Abstract

Group-living primates experience the benefits and costs associated with sociality, including an elevated risk of parasite transmission. However, the relative influence of group type (i.e., social structure), group size, and habitat disturbance on parasitic infection remains unclear, particularly in Southeast Asian primates. In this study, the abundance of intestinal parasites in proboscis monkeys (*Nasalis larvatus*) inhabiting a riverine forest along the Menanggul River, Sabah, Malaysian Borneo, was investigated. Fecal samples (n = 160) were collected from one-male-multifemale and all-male groups in areas with varying levels of anthropogenic disturbance, with efforts made to ensure that each sample originated from a different individual. In addition, the effects of group type, group size, and sampling location on parasite abundance were evaluated using fecal egg counts and Bayesian models. Three dominant parasite species groups (*Trichuris* sp., *Strongyloides fuelleborni*, and *Oesophagostomum aculeatum*) with an overall infection prevalence of 81.25% were identified. Results showed that group type did not significantly affect parasite abundance. However, group size showed a positive correlation with the abundance of *Trichuris* sp. and a negative correlation with *S. fuelleborni* and *O. aculeatum*. In addition, our models revealed that the infection load of *Trichuris* sp. decreased with increasing distance from the river mouth, which was used as a proxy for a disturbance gradient, whereas *O. aculeatum* exhibited higher infection load at greater distances, indicating lower prevalence in more disturbed downstream areas. Thus, parasite abundance in proboscis monkeys may be shaped by social and environmental factors, with taxa-specific responses likely reflecting differences in environmental persistence and transmission ecology.

## Introduction

Highly social animals such as primates exhibit complex group dynamics that influence their health, behavior, and interactions with pathogens (Kappeler et al., 2015). Although group-living promotes social support and cooperative behaviors that can mitigate some health risks (Silk, 2007), it also imposes costs, including an increased potential for pathogen transmission and heightened susceptibility to disease caused by social stress (Altizer et al., 2003). Thus, the relationship between sociality and disease risk is complicated. Factors such as group structure, contact patterns, and social hierarchies all play key roles in determining infection dynamics (Nunn et al., 2015). Notably, this relationship is bidirectional in primates. While social factors such as group size and composition influence parasite transmission, parasite infection can also feed back to shape host social behavior. Consequently, infection prevalence and intensity can affect social interaction and group dynamics (Vitone et al., 2004; Nunn, 2012; Rushmore et al., 2017).

Apart from social dynamics, anthropogenic environmental changes, such as deforestation and provisioning, can also influence parasite transmission by altering host movement patterns, population densities, and contact rates (Gillespie and Chapman, 2006; Becker et al., 2018). These ecological modifications can create conditions that either facilitate or hinder parasite spread, depending on the nature and extent of human-induced disturbances. Therefore, understanding the mechanism by which environmental, social, and ecological factors contribute to parasitic infections in social animals is crucial for developing effective conservation and management strategies (Chapman et al., 2005; Gillespie and Chapman, 2006).

On the contrary, research on parasitic infection and sociality in wild primates has yielded inconsistent findings, while the effects of anthropogenic environmental changes on parasitic infections in primates have been relatively consistent (Gillespie and Chapman, 2006). The relationship between parasite load and group size is complex, as it depends on various factors, such as host susceptibility and environmental conditions (e.g. Chapman et al., 2009; Klaus et al., 2018). Although larger groups might increase the likelihood of infection because of higher social interactions, group mobility and seasonal changes may reduce the risk of parasite transmission (e.g., increased precipitation during the rainy season washing away feces and parasites), thereby complicating the host–parasite dynamics (Nguyen et al., 2015; Radespiel et al., 2015; Knauf and Jones-Engel, 2020).

Although knowledge of parasitism in wild primates is expanding, it remains limited, particularly in Southeast Asia (Hopkins and Nunn, 2007; Cooper and Nunn, 2013). In Borneo, which is a highly biodiverse island in Southeast Asia, numerous endemic species are experiencing population declines (Wilcove et al., 2013), and the large arboreal proboscis monkey (*Nasalis larvatus*) inhabiting mangroves, peat swamps, and riverine forests is no exception to this trend (Lhota et al., 2019; Bernard et al., 2025). Several studies have provided insights into primate–parasite interactions in Sabah, including the proboscis monkey. This primate has unique parasite associations (Hasegawa et al., 2020) and infection dynamics that reflect its specialized ecological adaptations (Klaus et al., 2017) and interactions with sympatric primates (Frias et al., 2019; Frias et al., 2021). Previous studies have reported a high prevalence of helminth eggs in fecal samples, identifying five major parasite groups, with co-infections being common (Salgado-Lynn, 2010; Klaus et al., 2017; Frias et al., 2021). In addition, parasite species richness has been reported to be significantly higher in semi-wild populations than in wild ones. This increase likely results from the high levels of human disturbance, which can alter the mechanism by which parasites are transmitted. By contrast, group type and group size have been reported not to significantly affect parasite richness (Klaus et al., 2018). Despite the increasing knowledge of parasitic infections in proboscis monkeys, the fine-scale variation in infection patterns –across individuals or groups inhabiting distinct microhabitats and experiencing different levels of anthropogenic disturbance– remains unclear.

In this study, the abundance of intestinal parasites in proboscis monkeys was investigated, with particular emphasis on the effects of group type, group size, and habitat disturbance. Fecal samples collected over a 1-year period were analyzed to identify the parasites present and to examine the mechanism by which group size and social structure (i.e., group type: one-male-multifemale or all-male groups) influence parasite loads in a forest subjected to varying degrees of anthropogenic disturbance. Considering that proboscis monkeys exhibit seasonal variation in foraging behavior, ranging patterns, and intergroup spacing at our study site (Matsuda et al., 2009a; Matsuda et al., 2009b; Matsuda et al., 2010a), a year-round sampling approach may provide a comprehensive assessment of parasite dynamics. Thus, extended sampling could reveal a positive correlation between group size and parasitic infection, which is contrary to previous findings (Klaus et al., 2018). In addition, the study forest along the tributary river exhibits varying degrees of anthropogenic disturbance between the upstream and downstream areas. In particular, downstream areas are more strongly influenced by tourism-related boat traffic, human activity, and proximity to oil palm plantations, whereas upstream areas retain relatively higher plant diversity (Jose et al., 2024). The impact of habitat disturbance on parasite abundance can be assessed by comparing the fecal samples collected from individuals across these areas. Given that research in our study area has documented alterations in the gut microbiota of proboscis monkeys living in disturbed versus undisturbed areas (Jose et al., 2024), we hypothesize that parasite abundance would also differ between these environments because of human-induced disturbances.

## Methods

### Study site and subjects

This study was conducted in a riverine forest along the Menanggul River, a tributary of the Kinabatangan River, in Sabah, Malaysian Borneo (118°30′ E, 5°30′ N). The study site has a mean daily temperature range of 24°C–30°C and an annual average precipitation of 2474 mm (Matsuda et al., 2019). The river exhibits daily water-level fluctuations of approximately 1 m, while seasonal inundation results in an average increase exceeding 3 m (Matsuda et al., 2010b). The study area spans 4 km along the Menanggul River and supports a minimum population of 200 proboscis monkeys, forming a multilevel society composed of one-male-multifemale groups and all-male groups, which frequently assemble in the riverside (Matsuda et al., 2024). The riparian zone primarily consists of secondary forest in the southern portion, while much of the northern portion has been converted to oil palm plantations, except for a narrow, protected riparian strip. Accordingly, in this tributary, forests extending upstream of the river mouth exhibit a higher plant diversity (Jose et al., 2024). The proboscis monkeys in the study area are well habituated to human presence, as the Menanggul River has been a major tourist attraction for over a decade and serves as a frequent route for boats and observers.

### Sample collection

Between June 2015 and April 2016, boat surveys were conducted in the late afternoon on approximately 2–3 days per week to locate proboscis monkeys, as they typically return to riverside trees to sleep (Matsuda et al., 2011), and to record group composition and the GPS coordinates of sleeping locations. The following morning, before the monkeys awakened, these sleeping locations were revisited. Given that proboscis monkeys often defecate just before moving into the forest, the area around their sleeping sites was carefully searched for fresh feces after the monkeys had departed their sleeping trees. Multiple one-male-multifemale groups often remained close to one another along the river, making it challenging to determine to which group the feces found on the ground belonged. Therefore, we inferred the group associated with the feces based on the location of the feces and the group observed in the previous day’s survey. Only fecal samples presumed to originate from adult individuals were included, based on fecal size, which allows for differentiation between adults and immature individuals despite the challenges in distinguishing feces from neighboring groups (Matsuda et al., 2014). Consequently, between June 2015 and April 2016, 160 fecal samples were collected. Individual identity was subsequently confirmed through genetic analyses, which verified that the samples originated from genetically distinct individuals (Matsuda et al., 2024). At least nine groups were identified in the study area, although the number of groups these fecal samples were collected from remains unclear, as group identification was not complete at the time the samples were collected (Matsuda et al., 2024). The fecal samples were preserved in tubes with 70% ethanol.

### Parasite detection and identification

A modified protocol was used to concentrate parasite eggs via formalin-ethyl acetate sedimentation, and fecal samples were analyzed using the sequential sedimentation–flotation method (Frias et al., 2021). The final pellet was resuspended in saline, and 1 mL of aliquot was transferred into a vial and placed on a magnetic stirrer to ensure thorough homogenization during analysis. In estimating the number of eggs per gram of feces (EPG) in each sample, an aliquot was collected from the homogeneous suspension and placed in a McMaster counting chamber for examination at 100× magnification. EPG was calculated on the basis of the average of five replicate counts of parasite eggs observed within the chamber’s grid, using the known weight of fecal sediment in a 0.15 mL volume. After quantifying the eggs, each sample was centrifuged again; the supernatant was discarded, and the concentrated pellet was resuspended in Sheather’s solution with a specific gravity of 1.27. Two slides were examined to reduce the likelihood of overlooking less abundant helminth eggs. Parasite identification was performed using standard identification keys (Modrý et al., 2018).

### Parasite prevalence and abundance

The prevalence of a given parasite was calculated as the percentage of hosts infected by that parasite. Parasite abundance was estimated for each sample using the EPG rounded to the nearest integer as a proxy measure, following the terminology of Bush et al. (1997).

### Factors influencing parasite abundance

A Bayesian model was developed using the “brms” package (Bürkner, 2018) to evaluate the mechanism by which group type (one-male-multifemale or all-male groups), group size, and fecal sampling locations affect parasite abundance. Sampling locations were represented by distance from the river mouth to upstream locations, which served as a proxy for the disturbance gradient along the river system. In our analysis, parasite abundance was used as a dependent variable and modelled using a Poisson distribution with a log link. Group type was incorporated as a group-level effect, whereas group size and the distance of the sampling location to the river mouth were included as continuous variables. All models were controlled for host group identity. Distinct groups were not consistently identifiable throughout the study period. Therefore, we assigned samples to groups based on their spatial and temporal proximity along the river. This approach was guided by observations: the mean distance between consecutive sleeping sites for each group was approximately 350 m in the study area (Matsuda et al., 2024). In addition, changes in food availability over time can influence spatial patterns along the river (Matsuda et al., 2010a). Based on these criteria, two samples collected within 350 meters of each other and during the same month were classified as originating from the same group. Groups with fewer than three observations were excluded from the analysis.

Weakly informative priors were assigned to population-level effects, following a Student’s t-distribution. The posterior distributions of the model parameters were estimated using Markov chain Monte Carlo methods. For this purpose, four chains were constructed, each consisting of 2000 steps, with 1000-step warm-up periods. Consequently, a total of 4000 steps were retained to estimate the posterior distributions. The independent continuous variables were scaled to aid model fitting and coefficient interpretation. Chain convergence was assessed visually, and the potential scale reduction factor (Rhat) values were examined to ensure they were less than or equal to 1 for all parameters (Bürkner, 2018). This modeling process was repeated for the three parasites identified within the host population.

## Results

### Parasite prevalence and abundance

Parasites were detected in 130 of the 160 analyzed samples, yielding an overall parasite prevalence of 81.25%. The eggs of *Trichuris* sp. (78.12%, n = 125), *Strongyloides fuelleborni* (10.62%, n = 17), *Oesophagostomum aculeatum* (5%, n = 8), *Enterobius* (*Colobenterobius*) *serratus* (1.25%, n = 2), and *Spirurida* sp. (0.63%, n = 1) were identified. The mean abundance of the top three prevalent parasites was recorded at 168.2 EPG (range: 0–984), 9.41 EPG (range: 0–650), and 2.83 EPG (range: 0–109), respectively.

### Factors influencing parasite abundance

Of the 160 fecal samples examined, 124 were included in the Bayesian analyses after excluding samples from groups with fewer than three observations, as these could not be reliably incorporated into the group-level models. The type of host group did not have a statistically significant effect on parasite abundance across any of the parasite groups (Fig. 1A). However, host group size emerged as an important predictor of parasite abundance, with varying effects across parasite groups. The abundance of *Trichuris* sp. was positively correlated with group size, whereas that of *S. fuelleborni* and *O. aculeatum* was negatively correlated with group size (Fig. 1B). In addition, our models indicated that the distance from the river mouth to upstream locations, used as a proxy for a disturbance gradient, markedly influenced the parasite abundance for *Trichuris* sp. and *O. aculeatum*. In particular, as the distance from the river increased, the abundance of *Trichuris* sp. decreased, whereas *O. aculeatum* exhibited an opposite trend, with higher counts linked to greater distances from the river mouth (Fig. 1C). A summary of the results from the models is provided in Table 1.

**Fig. 1.**
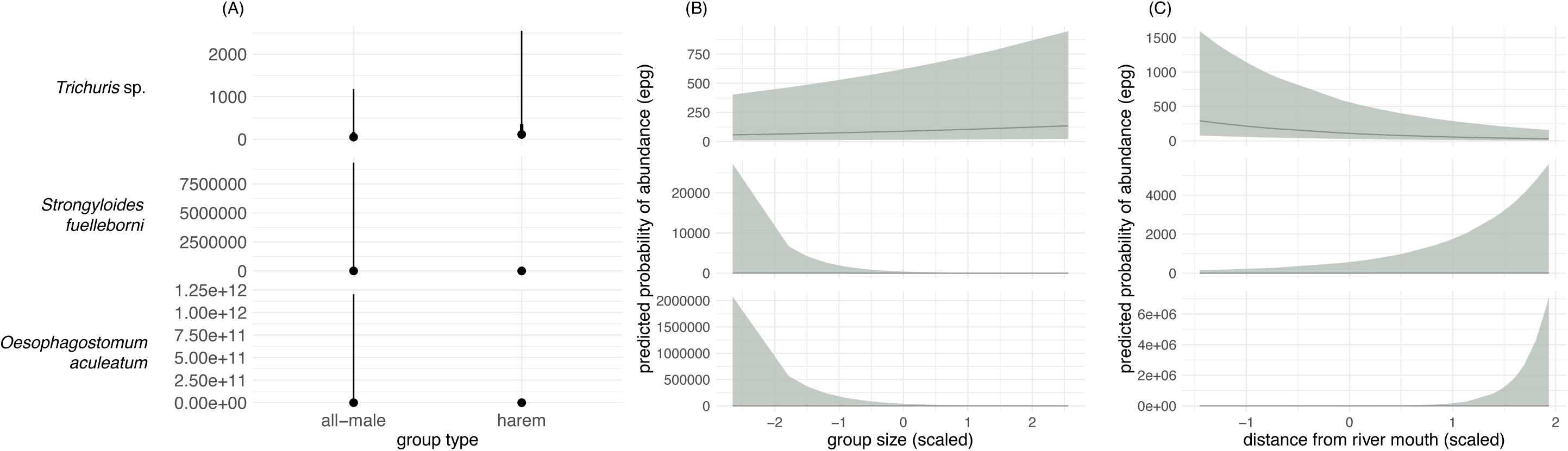
Posterior distributions of the differences in parasite abundance for *Trichuris* sp. (top), *S. fuelleborni* (middle), and *O. aculeatum* (bottom) in relation to host group type (A). The green shaded areas represent the posterior distributions, with the black dot indicating the median estimate and the black horizontal line shown in boldface representing the 99% credible interval. The vertical dashed line at zero represents no difference in infection rates among the groups. The predicted probability of abundance for the same parasite groups in relation to host group size (B) and sampling location represented by the distance from the river mouth (used as a proxy for disturbance gradient) (C). Credible intervals are set at 89%.

**Table 1.** Summary of the fitted models.

|  | Parameter | Est. | Est. error | 95% CI<br>(Lower) | 95% CI<br>(Upper) | Rhat | Bulk ESS | Tail ESS |
| --- | --- | --- | --- | --- | --- | --- | --- | --- |
| *Trichuris* sp. | *Random Effects* |  |  |  |  |  |  |  |
|  | sd (Intercept) | 0.92 | 0.22 | 0.6 | 1.44 | 1 | 1600 | 2336 |
|  | *Fixed Effects* |  |  |  |  |  |  |  |
|  | Intercept | 4.32 | 0.51 | 3.29 | 5.32 | 1 | 3515 | 4008 |
|  | group type | 0.81 | 0.55 | -0.27 | 1.93 | 1 | 3151 | 3511 |
|  | group size (scaled) | 0.16 | 0.01 | 0.14 | 0.19 | 1 | 6937 | 4494 |
|  | distance to river (scaled) | -0.7 | 0.14 | -0.97 | -0.44 | 1 | 2949 | 3362 |
| *S. fuelleborni* | *Random Effects* |  |  |  |  |  |  |  |
|  | sd (Intercept) | 5.74 | 1.67 | 3.37 | 9.71 | 1 | 2691 | 3424 |
|  | *Fixed Effects* |  |  |  |  |  |  |  |
|  | Intercept | -1.68 | 3.25 | -8.21 | 4.77 | 1 | 3101 | 3670 |
|  | group type | -1.09 | 3.7 | -8.69 | 5.88 | 1 | 2916 | 3602 |
|  | group size (scaled) | -1.62 | 0.12 | -1.86 | -1.38 | 1 | 9301 | 5682 |
|  | distance to river (scaled) | 1.03 | 0.71 | -0.35 | 2.44 | 1 | 5057 | 4881 |
| *O. aculeatum* | *Random Effects* |  |  |  |  |  |  |  |
|  | sd (Intercept) | 10.54 | 3.96 | 5 | 20.28 | 1 | 2116 | 3425 |
|  | *Fixed Effects* |  |  |  |  |  |  |  |
|  | Intercept | -5.12 | 6.63 | -18.87 | 7.67 | 1 | 3150 | 3920 |
|  | group type | -0.47 | 7.58 | -16.08 | 14.32 | 1 | 2693 | 3620 |
|  | group size (scaled) | -1.48 | 0.19 | -1.87 | -1.11 | 1 | 7942 | 5445 |
|  | distance to river (scaled) | 3.46 | 1.4 | 0.71 | 6.27 | 1 | 3953 | 4708 |

## Discussion

In this study, the relationship among social structure (i.e., group type: one-male-multifemale and all-male groups), the degree of anthropogenic habitat disturbances within a 4-km stretch along the river, and parasite load in free-ranging proboscis monkeys was examined. A high prevalence of intestinal parasites was observed (81.25%), which is consistent with the previous prevalences recorded for proboscis monkeys along the Kinabatangan River in Sabah [i.e., 96.6% in Salgado-Lynn (2010), 92.3% in Klaus et al. (2017), and 80.2% in Frias et al. (2021)] and in Bako National Park in Sarawak [i.e., 95.4% in Adrus et al. (2019)]. The predominant parasite groups in this study included *Trichuris* sp., *Strongyloides fuelleborni*, and *Oesophagostomum aculeatum*, which have also been identified as dominant in other fecal analyses of proboscis monkeys, in wild and captive settings (Table 2), reinforcing a consistent pattern of parasitic infections across different populations. Nonetheless, given the substantial differences in the diets of proboscis monkeys between riverine and mangrove forests (Matsuda et al., 2019), habitat-linked variation in food use may also influence parasite exposure. Because habitat type determines not only the food resources available to proboscis monkeys but also forest height and frequency of ground use or sediment contact during foraging (Feilen and Marshall, 2014, 2020; Atmoko et al., 2021), these ecological differences may affect transmission dynamics of environmentally transmitted parasites. Thus, whether similar trends in parasite prevalence are observed among proboscis monkeys across different habitat types on the island of Borneo should be examined in the future.

**Table 2.**
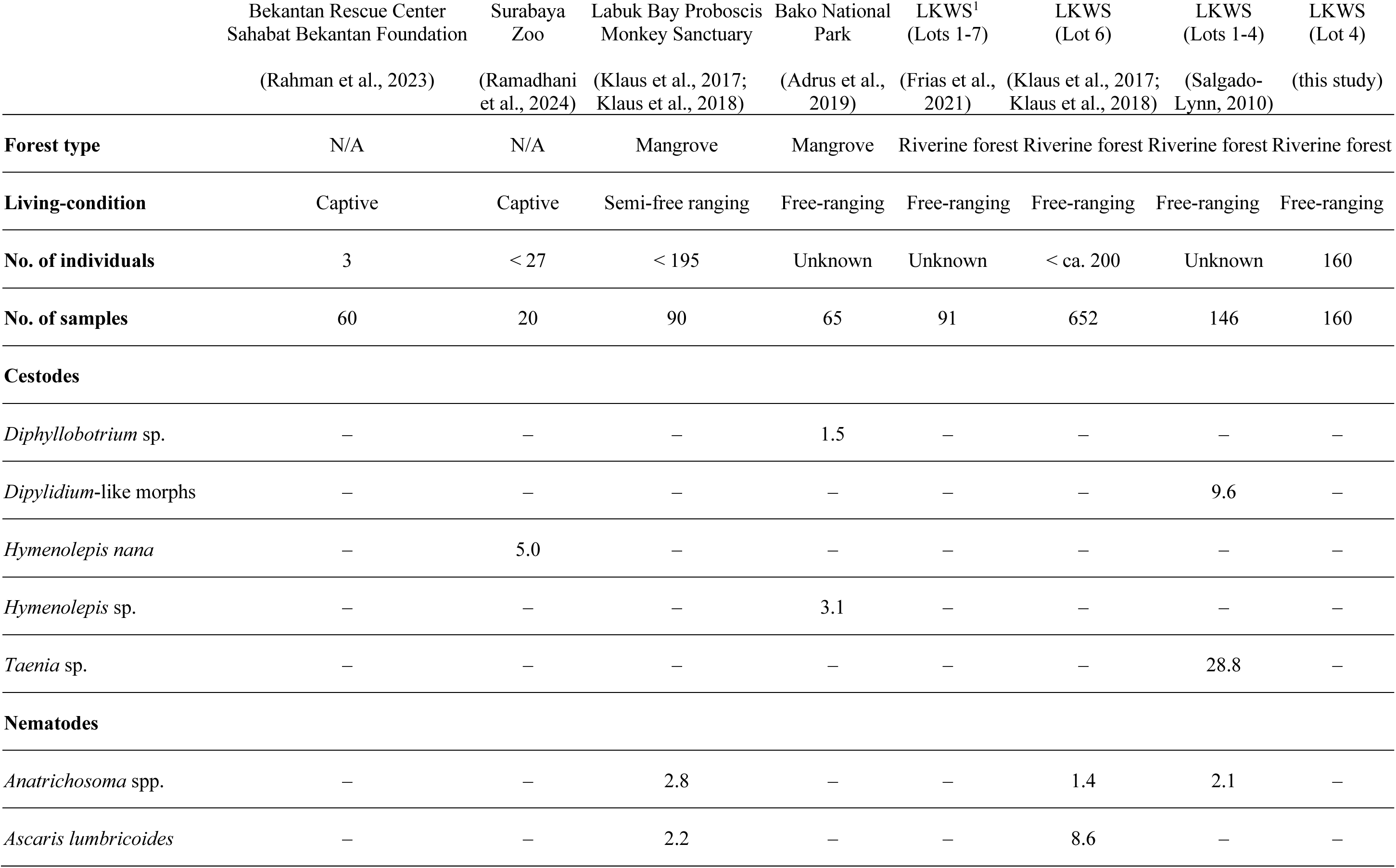
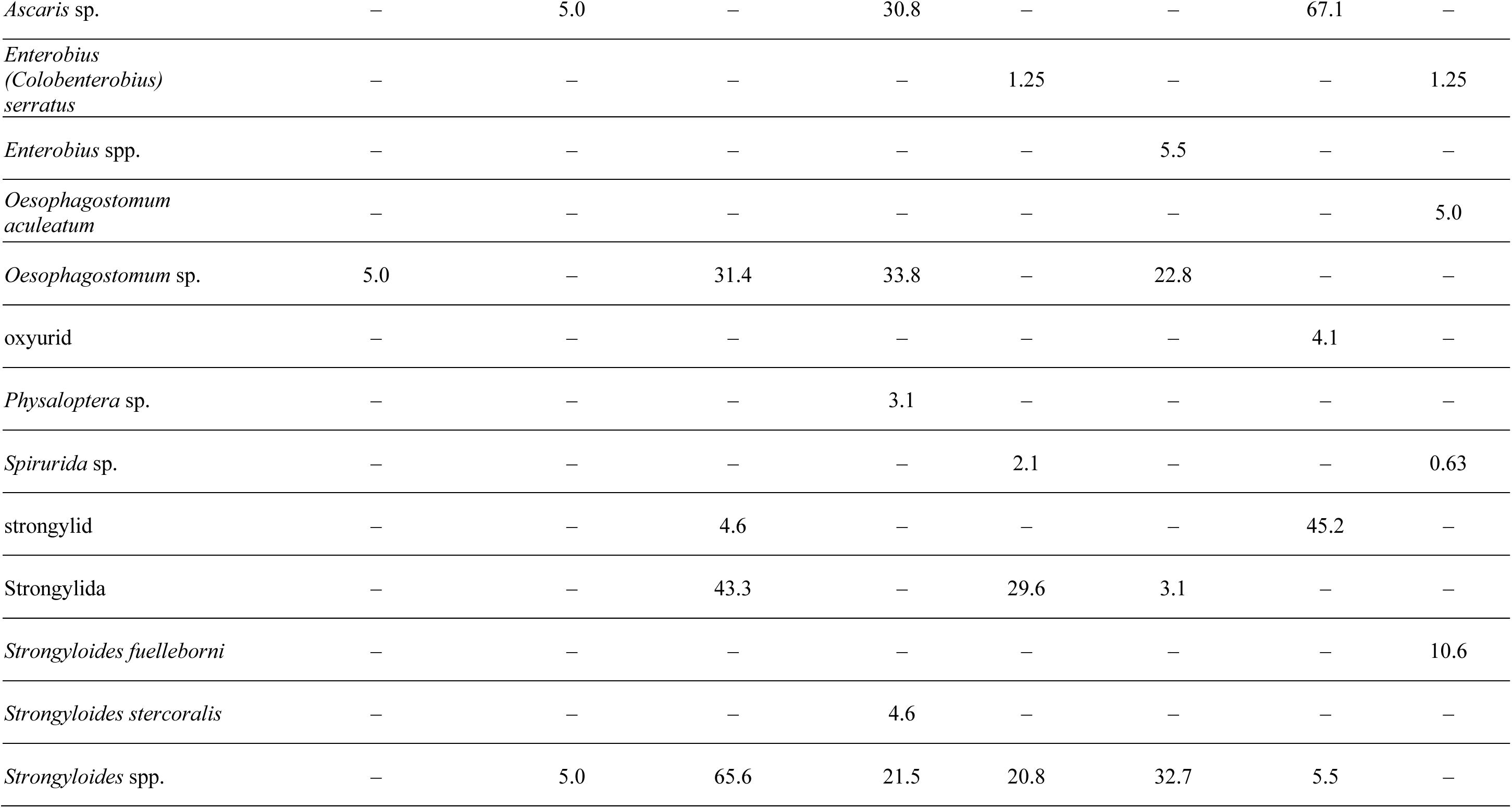
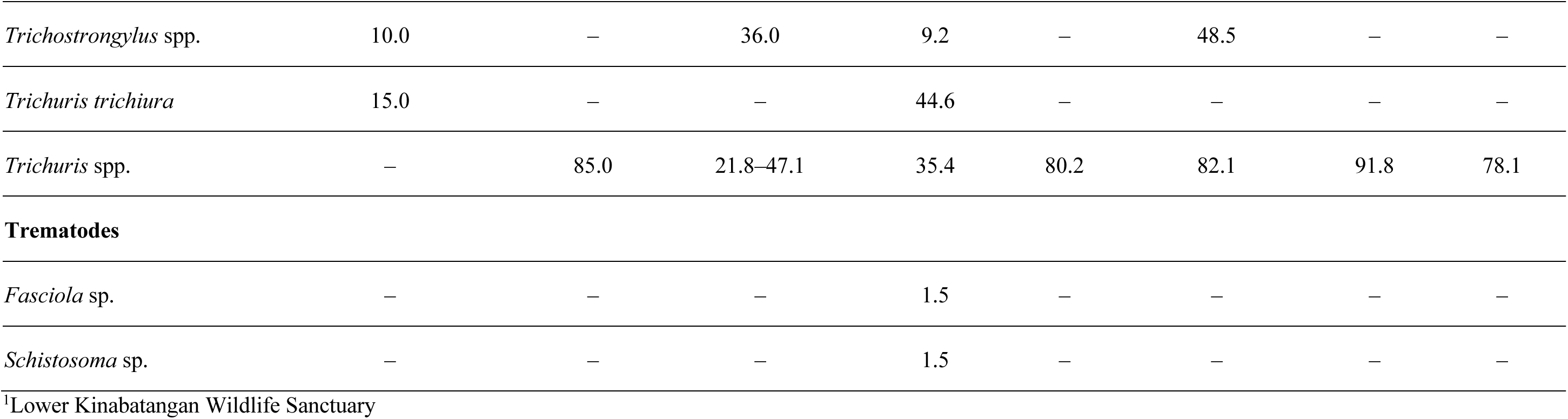
Comparison between the parasite species detected and parasite prevalence (%) in fecal samples of proboscis monkeys in each study.

However, no significant association was found between parasite abundance and group type (i.e., one-male-multifemale and all-male groups) in proboscis monkeys in this study, consistent with previous findings (Klaus et al., 2018), suggesting that sex differences may have a limited influence on parasitic infection. This interpretation is further supported by a study conducted at the same site that examined gut microbial composition in fecal samples and found no significant sex differences (Jose et al., 2024). Given that gut microbiota, such as intestinal parasites, would be transmitted through direct or indirect social contact and shared use of space and food resources, the absence of sex-specific patterns in both types of organisms indicates that behavioral and spatial overlap between males and females minimizes sex-related divergence in microbial and parasite exposure.

Contrary to the findings of Klaus et al. (2018), our study revealed that group size might be a potential factor influencing parasite abundance for a particular parasite species, namely, *Trichuris* sp. This finding supports the hypothesis that larger groups may exhibit higher infection rates, probably because of increased contact rates and overlapping space use among individuals (Snaith et al., 2008; Rifkin et al., 2012; Kappeler et al., 2015). Although the exact mechanism remains unclear, a comparable trend was observed in the gut microbial diversity in the same population. A larger group size was associated with greater alpha diversity (Jose et al., 2024), suggesting that higher host density may increase exposure to intestinal parasites and microbes. This association may reflect the increased direct interactions among individuals and result from enhanced environmental contamination in frequently reused areas. Given that all the parasites identified in this study could be transmitted via fecal-oral routes, the accumulation of feces in shared sleeping or foraging sites could be a key factor driving infection. Limited territoriality and habitual reuse of sleeping trees by proboscis monkeys –often shared across one-male-multifemale and all-male groups in multilevel societies– likely facilitates such transmission (Matsuda et al., 2024). Furthermore, the differences in habitat structure and spatial dynamics among the study sites may explain the contrasting results reported by Klaus et al. (2018). They focused on groups inhabiting the main channel of the Kinabatangan River, where the river’s width (>200 m) limits cross-group movement and shared space use. By contrast, our study site along narrow tributaries (15–20 m wide) may allow more frequent crossings and higher spatial overlap among groups (Matsuda et al., 2008; Matsuda et al., 2010a), thereby promoting indirect transmission through contaminated environments.

On the contrary, *S. fuelleborni* and *O. aculeatum* were negatively correlated with group size because of their differing ecological tolerances. *S. fuelleborni*, which is a parasitic nematode, may be more influenced by habitat disturbance or seasonal changes, with larger groups potentially reducing parasite exposure through increased host mobility and spatial separation from contaminated areas (Gillespie and Chapman, 2006; Radespiel et al., 2015). Similarly, *O. aculeatum* may require more stable environmental conditions for successful transmission, as the development and survival of free-living stages in *Oesophagostomum* spp. are influenced by factors such as temperature and related habitat conditions (Sengupta et al., 2016; Senanayake et al., 2025). The relatively undisturbed and pristine upstream habitats may therefore provide better conditions for transmission compared with downstream areas, whereas larger groups in disturbed downstream areas could hinder transmission (Kappeler et al., 2015). More broadly, these contrasting patterns may reflect a trade-off between contamination density and spatial avoidance. Larger groups may increase fecal contamination at frequently reused sites, favoring parasites with environmentally resistant stages such as *Trichuris*. Conversely, increased host mobility and shorter residence time at specific patches may reduce exposure to parasites with more fragile free-living stages, such as *S. fuelleborni* and *O. aculeatum*. These findings indicate that host social dynamics and environmental factors must be considered, as species-specific responses shape infection patterns (Rifkin et al., 2012; Nguyen et al., 2015).

Our findings revealed a more complex relationship between parasite abundance and habitat disturbance than initially anticipated. Contrary to expectations that anthropogenic habitat disturbance would universally elevate parasite loads (Gillespie and Chapman, 2006), we observed that the impact of disturbance varied across different parasite taxa. In particular, the abundance of *Trichuris* sp. decreased with increasing distance from the river mouth, indicating that higher parasite loads occur in downstream areas with more intense human activity and tourism. Conversely, *O. aculeatum* displayed the opposite trend, with greater abundance in less disturbed upstream forests. These contrasting patterns indicate that different parasite species may respond to habitat disturbance through distinct ecological mechanisms. However, we acknowledge that distance from the river mouth serves as a proxy that likely integrates multiple environmental gradients beyond anthropogenic disturbance alone. These may include natural variation in soil composition, flooding regimes, vegetation structure, and microclimatic stability, all of which could influence parasite spatial responses observed between *Trichuris* sp. and *O. aculeatum*.

This interpretation would be consistent with differences in environmental persistence between parasite taxa, as *Trichuris* eggs are relatively resistant, whereas *O. aculeatum* may depend more strongly on stable microenvironmental conditions for successful development and transmission. Considering the elevated *Trichuris* sp. abundance in downstream areas, more disturbed habitats may not be attributable to the increased host density or spatial overlap, as proboscis monkeys in these areas do not necessarily aggregate at higher densities compared with the upstream areas. Aggregation likely occurs in upstream areas, where the river is narrower, providing a safer environment for proboscis monkeys because they can escape terrestrial predators, such as clouded leopards, by jumping to the opposite bank (Matsuda et al., 2010a). Given its greater safety from predators, this environment likely facilitates aggregation in upstream areas, making them a more favorable location for proboscis monkey groups to congregate. This interpretation is consistent with the species’ highly resilient egg morphology, which supports environmental persistence even under degraded conditions (Gillespie and Chapman, 2006; Snaith et al., 2008). Morphological robustness may enable *Trichuris* sp. to maintain transmission success across a range of habitat types, including more disturbed downstream areas. By contrast, successful transmission of *O. aculeatum* requires embryonation, hatching, and survival of infective larval stages in the environment, processes that are influenced by environmental conditions such as temperature and moisture (Rose and Small, 1980; Radespiel et al., 2015; Sengupta et al., 2016; Senanayake et al., 2025). Accordingly, the relatively pristine conditions in upstream sites likely provide a favorable microenvironment for their development.

Interestingly, these spatial patterns in parasite abundance are contrary to prior findings from the same population, which identified clear site-based differences in gut microbial composition, with lower microbial diversity in the downstream areas (Jose et al., 2024). Gut microbiota and intestinal parasites share common transmission pathways, including social contact and environmental exposure. However, their ecological sensitivities appear to diverge. In general, gut microbiota are more responsive to dietary shifts and subtle ecological changes (e.g., Amato et al., 2013; Hayakawa et al., 2018; Lee et al., 2019; Barelli et al., 2020). By contrast, parasite abundance in our study is influenced by a combination of host social dynamics (e.g., group size and space use) and species-specific environmental tolerances. This distinction emphasizes the importance of adopting a taxon-specific approach when interpreting the host–environment–parasite relationship. Furthermore, it indicates that parasites and microbiota may differentially influence the ecological and health dynamics of primates (Rifkin et al., 2012; Nguyen et al., 2015).

These contrasting patterns between gut microbiota diversity and *Trichuris* abundance may reflect differences in the ecological processes governing their assembly. Gut microbiota are strongly shaped by dietary and environmental inputs, which can be altered by habitat disturbance (Amato et al., 2013; Hayakawa et al., 2018). In contrast, helminth infections depend more directly on encounters with infective stages in the environment and patterns of fecal contamination (Nunn et al., 2011). Parasites with environmentally resistant eggs, such as *Trichuris*, may therefore persist and accumulate even in disturbed habitats where microbial exposure diversity declines. Disturbed environments may thus simultaneously reduce opportunities for microbial acquisition while maintaining or even increasing exposure to environmentally persistent helminths.

Understanding the interplay among social organization, habitat characteristics, and parasitic infection in wild primates deepens our ecological insights. It informs broader questions in evolutionary biology and conservation. The findings of this study underscore the complex, context-dependent nature of host–parasite dynamics. They highlight how intrinsic factors such as group size and behavior can mediate exposure risks, even in the face of external environmental change. Given the escalating habitat disturbance, a nuanced understanding is crucial for predicting disease risks in wildlife populations and for designing conservation strategies that account for social and ecological variability. Integrative approaches that consider behavioral ecology alongside environmental gradients may also be essential for safeguarding the health and persistence of socially complex yet increasingly vulnerable species such as the proboscis monkey.

Finally, the helminths detected in this study are of potential zoonotic relevance, especially in ecotourism landscapes such as the Menanggul River, where frequent contact occurs among humans, livestock, and wild primates. The genus *Trichuris* may include species closely related to *T. trichiura*, a well-known human pathogen that can infect non-human primates (Rivero et al., 2020). Likewise, *S. fuelleborni* and *O. aculeatum* have been reported in both human and non-human hosts in Southeast Asia (Yalcindag et al., 2021; Chan et al., 2023). However, because parasite identification in the present study was based solely on egg morphology, neither host specificity nor actual zoonotic transmission risk can be confirmed at this stage. Molecular characterisation will therefore be required to determine whether the detected parasites belong to human-infective lineages. Nevertheless, the presence of these helminths in areas accessible to humans highlights the need for further investigation of their transmission dynamics and potential public health relevance under increasing anthropogenic pressure (Gillespie and Chapman, 2006).

## Acknowledgements

We express our sincere thanks to the Sabah Biodiversity Centre and the Sabah Wildlife Department for granting permission to carry out this research. We are also deeply indebted to the Sabah Forestry Department for facilitating the use of its field facilities. We particularly thank the research assistants, Asnih Binti Etin and Jasrudy Bin Mandu, for their support in the field. Finally, we are grateful to the reviewers for their fruitful comments that improved this manuscript.

## Author contribution

MNF: conceptualization, writing—original draft, methodology; LF: conceptualization, writing—review and editing, methodology, visualization, formal analysis, investigation; EZ: conceptualization, methodology; PL: resources, writing—review and editing; JT: resources, writing—review and editing; HB: resources, writing—review and editing; VK: resources, writing—review and editing; IM: conceptualization, data curation, funding acquisition, investigation, methodology, project administration, resources, supervision, validation, visualization, writing—original draft. All authors (except deceased EZ) gave final approval for publication and agreed to be held accountable for the work performed therein.

## Data availability

All data generated or analysed during this study are included in this published article.

## Declarations

### Ethics approval

The animal study was reviewed and approved by the Sabah Biodiversity Centre and the Sabah Wildlife Department [JKM/MBS.1000-2/2 JLD.3 (120)].

### Generative AI and AI-assisted technologies in the writing process

During the preparation of this manuscript, the authors used ChatGPT-5.2 to refine the clarity and logical flow of the text. The authors carefully reviewed, corrected, and approved all content generated, and take full responsibility for the final published version.

## References

Adrus, M., Zainuddin, R., Ahmad Khairi, N.H., Ahamad, M., Abdullah, M.T., 2019. Helminth parasites occurrence in wild proboscis monkeys (*Nasalis larvatus*), endemic primates to Borneo Island. J Med Primatol 48, 357–363. doi: 10.1111/jmp.12437

Altizer, S., Nunn, C.L., Thrall, P.H., Gittleman, J.L., Antonovics, J., Cunningham, A.A., Dobson, A.P., Ezenwa, V., Jones, K.E., Pedersen, A.B., Poss, M., Pulliam, J.R.C., 2003. Social Organization and Parasite Risk in Mammals: Integrating Theory and Empirical Studies. Annual Review of Ecology, Evolution, and Systematics 34, 517–547. doi: 10.1146/annurev.ecolsys.34.030102.151725

Amato, K.R., Yeoman, C.J., Kent, A., Righini, N., Carbonero, F., Estrada, A., Gaskins, H.R., Stumpf, R.M., Yildirim, S., Torralba, M., Gillis, M., Wilson, B.A., Nelson, K.E., White, B.A., Leigh, S.R., 2013. Habitat degradation impacts black howler monkey (*Alouatta pigra*) gastrointestinal microbiomes. ISME J. 7, 1344–1353. doi: 10.1038/ismej.2013.16

Atmoko, T., Mardiastuti, A., Bismark, M., Prasetyo, L.B., Iskandar, E., 2021. Land cover and Proboscis monkey habitats in Berau Delta, East Kalimantan. IOP Conference Series: Earth and Environmental Science 739. doi: 10.1088/1755-1315/739/1/012062

Barelli, C., Albanese, D., Stumpf, R.M., Asangba, A., Donati, C., Rovero, F., Hauffe, H.C., 2020. The gut microbiota communities of wild arboreal and ground-feeding tropical primates are affected differently by habitat disturbance. mSystems 5. doi: 10.1128/mSystems.00061-20

Becker, D.J., Streicker, D.G., Altizer, S., 2018. Using host species traits to understand the consequences of resource provisioning for host-parasite interactions. J Anim Ecol 87, 511–525. doi: 10.1111/1365-2656.12765

Bernard, H., Mohammad-Shom, S., Kulanthavelu, M., Sha, J.C.M., Malim, T.P., Abram, N.K., Matsuda, I., 2025. Monitoring the population and distribution of the proboscis monkey (*Nasalis larvatus*) in the Klias Peninsula, Sabah, Borneo, Malaysia: insights from an 18-year study. Primates 66, 277–294. doi: 10.1007/s10329-025-01183-7

Bürkner, P.-C., 2018. Advanced Bayesian Multilevel Modeling with the R Package brms. The R Journal 10, 395–411. doi:

Bush, A.O., Lafferty, K.D., Lotz, J.M., Shostak, A.W., 1997. Parasitology meets ecology on its own terms: Margolis et al. revisited. The Journal of Parasitology 83. doi: 10.2307/3284227

Chan, A.H.E., Kusolsuk, T., Watthanakulpanich, D., Pakdee, W., Doanh, P.N., Yasin, A.M., Dekumyoy, P., Thaenkham, U., 2023. Prevalence of *Strongyloides* in Southeast Asia: a systematic review and meta-analysis with implications for public health and sustainable control strategies. Infect Dis Poverty 12, 83. doi: 10.1186/s40249-023-01138-4

Chapman, A.C., Rothman, J.M., Hodder, S.A.M., 2009. Can parasite infections be a selective force influencing primate group size? A test with red colobus, in: Huffman, M.A., Chapman, C.A. (Eds.), Primate Parasite Ecology: The Dynamics and Study of Host-Parasite Relationships. Cambridge University Press, Cambridge, pp. 423–440.

Chapman, C.A., Gillespie, T.R., Goldberg, T.L., 2005. Primates and the Ecology of their Infectious Diseases: How will Anthropogenic Change Affect Host-Parasite Interactions? Evolutionary Anthropology: Issues, News, and Reviews 14, 134–144. doi: 10.1002/evan.20068

Cooper, N., Nunn, C.L., 2013. Identifying future zoonotic disease threats: Where are the gaps in our understanding of primate infectious diseases? Evol Med Public Health 2013, 27–36. doi: 10.1093/emph/eot001

Feilen, K.L., Marshall, A.J., 2014. Sleeping site selection by proboscis monkeys (*Nasalis larvatus*) in West Kalimantan, Indonesia. Am J Primatol 76, 1127–1139. doi: 10.1002/ajp.22298

Feilen, K.L., Marshall, A.J., 2020. Responses to spatial and temporal variation in food availability on the feeding ecology of proboscis monkeys (*Nasalis larvatus*) in West Kalimantan, Indonesia. Folia Primatol (Basel), 1–18. doi: 10.1159/000504362

Frias, L., Hasegawa, H., Chua, T.H., Sipangkui, S., Stark, D.J., Salgado-Lynn, M., Goossens, B., Keuk, K., Okamoto, M., MacIntosh, A.J.J., 2021. Parasite community structure in sympatric Bornean primates. International Journal for Parasitology. doi: 10.1016/j.ijpara.2021.03.003

Frias, L., Stark, D.J., Salgado Lynn, M., Nathan, S., Goossens, B., Okamoto, M., MacIntosh, A.J.J., 2019. Molecular characterization of nodule worm in a community of Bornean primates. Ecology and Evolution. doi: 10.1002/ece3.5022

Gillespie, T.R., Chapman, C.A., 2006. Prediction of parasite infection dynamics in primate metapopulations based on attributes of forest fragmentation. Conserv Biol 20, 441–448. doi: 10.1111/j.1523-1739.2006.00290.x

Hasegawa, H., Frias, L., Peter, S., Hasan, N.H., Stark, D.J., Lynn, M.S., Sipangkui, S., Goossens, B., Matsuura, K., Okamoto, M., Macintosh, A.J.J., 2020. First description of male worms of *Enterobius* (*Colobenterobius*) *serratus* (Nematoda: Oxyuridae), the pinworm parasite of proboscis monkeys. Zootaxa 4722, 287–294. doi: 10.11646/zootaxa.4722.3.6

Hayakawa, T., Nathan, S., Stark, D.J., Saldivar, D.A.R., Sipangkui, R., Goossens, B., Tuuga, A., Clauss, M., Sawada, A., Fukuda, S., Imai, H., Matsuda, I., 2018. First report of foregut microbial community in proboscis monkeys: are diverse forests a reservoir for diverse microbiomes? Environmental Microbiology Reports 10, 655–662. doi: 10.1111/1758-2229.12677

Hopkins, M.E., Nunn, C.L., 2007. A global gap analysis of infectious agents in wild primates. Diversity and Distributions 13, 561–572. doi: 10.1111/j.1472-4642.2007.00364.x

Jose, L., Lee, W., Hanya, G., Tuuga, A., Goossens, B., Tangah, J., Matsuda, I., Kumar, V.S., 2024. Gut microbial community in proboscis monkeys (*Nasalis larvatus*): implications for effects of geographical and social factors. Royal Society Open Science 11. doi: 10.1098/rsos.231756

Kappeler, P.M., Cremer, S., Nunn, C.L., 2015. Sociality and health: impacts of sociality on disease susceptibility and transmission in animal and human societies. Philos Trans R Soc Lond B Biol Sci 370. doi: 10.1098/rstb.2014.0116

Klaus, A., Strube, C., Roper, K.M., Radespiel, U., Schaarschmidt, F., Nathan, S., Goossens, B., Zimmermann, E., 2018. Fecal parasite risk in the endangered proboscis monkey is higher in an anthropogenically managed forest environment compared to a riparian rain forest in Sabah, Borneo. PLoS One 13, e0195584. doi: 10.1371/journal.pone.0195584

Klaus, A., Zimmermann, E., Röper, K.M., Radespiel, U., Nathan, S., Goossens, B., Strube, C., 2017. Co-infection patterns of intestinal parasites in arboreal primates (proboscis monkeys, Nasalis larvatus) in Borneo. International Journal for Parasitology: Parasites and Wildlife. doi: 10.1016/j.ijppaw.2017.09.005

Knauf, S., Jones-Engel, L., 2020. Neglected Diseases in Monkeys: From the Monkey-Human Interface to One Health. Springer Cham, Switzerland AG.

Lee, W., Hayakawa, T., Kiyono, M., Yamabata, N., Hanya, G., 2019. Gut microbiota composition of Japanese macaques associates with extent of human encroachment. Am J Primatol 81, e23072. doi: 10.1002/ajp.23072

Lhota, S., Sha, J.C.M., Bernard, H., Matsuda, I., 2019. Proboscis monkey conservation: beyond the science, in: Behie, A.M., Teichroeb, J.A., Malone, N.M. (Eds.), Primate Research and Conservation in the Anthropocene. Cambridge University Press, Cambridge pp. 182–196.

Matsuda, I., Kubo, T., Tuuga, A., Higashi, S., 2010a. A Bayesian analysis of the temporal change of local density of proboscis monkeys: implications for environmental effects on a multilevel society. American Journal of Physical Anthropology 142, 235–245. doi: 10.1002/ajpa.21218

Matsuda, I., Murai, T., Grueter, C.C., Tuuga, A., Goossens, B., Bernard, H., Yahya, N.K., Orozco-terWengel, P., Salgado-Lynn, M., 2024. The multilevel society of proboscis monkeys with a possible patrilineal basis. Behavioral Ecology and Sociobiology 78. doi: 10.1007/s00265-023-03419-2

Matsuda, I., Nakabayashi, M., Otani, Y., Yap, S.W., Tuuga, A., Wong, A., Bernard, H., Wich, S.A., Kubo, T., 2019. Comparison of plant diversity and phenology of riverine and mangrove forests with those of the dryland forest in Sabah, Borneo, Malaysia, in: Nowak, K., Barnett, A.A., Matsuda, I. (Eds.), Primates in Flooded Habitats: Ecology and Conservation. Cambridge University Press, Cambridge, pp. 15–28.

Matsuda, I., Tuuga, A., Akiyama, Y., Higashi, S., 2008. Selection of river crossing location and sleeping site by proboscis monkeys (*Nasalis larvatus*) in Sabah, Malaysia. American Journal of Primatology 70, 1097–1101. doi: 10.1002/ajp.20604

Matsuda, I., Tuuga, A., Bernard, H., 2011. Riverine refuging by proboscis monkeys (*Nasalis larvatus*) and sympatric primates: Implications for adaptive benefits of the riverine habitat. Mammalian Biology 76, 165–171. doi: 10.1016/j.mambio.2010.03.005

Matsuda, I., Tuuga, A., Hashimoto, C., Bernard, H., Yamagiwa, J., Fritz, J., Tsubokawa, K., Yayota, M., Murai, T., Iwata, Y., Clauss, M., 2014. Faecal particle size in free-ranging primates supports a ‘rumination’ strategy in the proboscis monkey (*Nasalis larvatus*). Oecologia 174, 1127–1137. doi: 10.1007/s00442-013-2863-9

Matsuda, I., Tuuga, A., Higashi, S., 2009a. The feeding ecology and activity budget of proboscis monkeys. American Journal of Primatology 71, 478–492. doi: 10.1002/ajp.20677

Matsuda, I., Tuuga, A., Higashi, S., 2009b. Ranging behavior of proboscis monkeys in a riverine forest with special reference to ranging in inland forest. International Journal of Primatology 30, 313–325. doi: 10.1007/s10764-009-9344-3

Matsuda, I., Tuuga, A., Higashi, S., 2010b. Effects of water level on sleeping-site selection and inter-group association in proboscis monkeys: why do they sleep alone inland on flooded days? Ecological Research 25, 475–482. doi: 10.1007/s11284-009-0677-3

Modrý, D., Pafcǒ, B., Petrzělková, K.J., Hasegawa, H., 2018. Parasites of Apes: an Atlas of Coproscopic Diagnostics. Chimaira, Frankfurt.

Nguyen, N., Fashing, P.J., Boyd, D.A., Barry, T.S., Burke, R.J., Goodale, C.B., Jones, S.C., Kerby, J.T., Kellogg, B.S., Lee, L.M., Miller, C.M., Nurmi, N.O., Ramsay, M.S., Reynolds, J.D., Stewart, K.M., Turner, T.J., Venkataraman, V.V., Knauf, Y., Roos, C., Knauf, S., 2015. Fitness impacts of tapeworm parasitism on wild gelada monkeys at Guassa, Ethiopia. Am J Primatol 77, 579–594. doi: 10.1002/ajp.22379

Nunn, C.L., 2012. Primate disease ecology in comparative and theoretical perspective. Am J Primatol 74, 497–509. doi: 10.1002/ajp.21986

Nunn, C.L., Jordan, F., McCabe, C.M., Verdolin, J.L., Fewell, J.H., 2015. Infectious disease and group size: more than just a numbers game. Philos Trans R Soc Lond B Biol Sci 370. doi: 10.1098/rstb.2014.0111

Nunn, C.L., Thrall, P.H., Leendertz, F.H., Boesch, C., 2011. The spread of fecally transmitted parasites in socially-structured populations. PLoS One 6, e21677. doi: 10.1371/journal.pone.0021677

Radespiel, U., Schaber, K., Kessler, S.E., Schaarschmidt, F., Strube, C., 2015. Variations in the excretion patterns of helminth eggs in two sympatric mouse lemur species (*Microcebus murinus* and *M. ravelobensis*) in northwestern Madagascar. Parasitol Res 114, 941–954. doi: 10.1007/s00436-014-4259-0

Rahman, F.G., Nurliani, A., Rezeki, A., Rusmiati, R., 2023. Identification of intestine parasite worms eggs in feces proboscis monkey (*Nasalis larvatus*). Proceeding of International Conference on Biology Education, Natural Science, and Technology 1, 541–549. doi:

Ramadhani, M., Nurhidayah, N., Kurniawati, D.A., Purmadi, M., Rafiq, F.A., Kusumaningtyas, E., Endrawati, D., Hidayatik, N., Prastiya, R.A., Mariya, S.S., Mufasirin, Hastutiek, P., 2024. Gastrointestinal helminths of captive proboscis monkey (*Nasalis larvatus*) in Surabaya zoo, Indonesia. J Med Primatol 53, e12719. doi: 10.1111/jmp.12719

Rifkin, J.L., Nunn, C.L., Garamszegi, L.Z., 2012. Do animals living in larger groups experience greater parasitism? A meta-analysis. Am Nat 180, 70–82. doi: 10.1086/666081

Rivero, J., Cutillas, C., Callejon, R., 2020. *Trichuris trichiura* (Linnaeus, 1771) From Human and Non-human Primates: Morphology, Biometry, Host Specificity, Molecular Characterization, and Phylogeny. Front Vet Sci 7, 626120. doi: 10.3389/fvets.2020.626120

Rose, J.H., Small, A.J., 1980. Observations on the development and survival of the free-living stages of Oesophagostomum dentatum both in their natural environments out-of-doors and under controlled conditions in the laboratory. Parasitology 81, 507–517. doi: 10.1017/s0031182000061898

Rushmore, J., Bisanzio, D., Gillespie, T.R., 2017. Making New Connections: Insights from Primate-Parasite Networks. Trends Parasitol 33, 547–560. doi: 10.1016/j.pt.2017.01.013

Salgado-Lynn, M., 2010. Primate viability in a fragmented landscape: genetic diversity and parasite burden of long-tailed macaques and proboscis monkeys in the Lower Kinabatangan Floodplain, Sabah, Malaysia. Cardiff University, Cardiff.

Senanayake, N.S., Boyle, L., O’Driscoll, K., Menant, O., Butler, F., 2025. Effects of season, age and parasite management practices on gastro - intestinal parasites in pigs kept outdoors in Ireland. Ir Vet J 78, 12. doi: 10.1186/s13620-025-00297-0

Sengupta, M.E., Thapa, S., Thamsborg, S.M., Mejer, H., 2016. Effect of vacuum packing and temperature on survival and hatching of strongyle eggs in faecal samples. Vet Parasitol 217, 21–24. doi: 10.1016/j.vetpar.2015.12.014

Silk, J.B., 2007. The adaptive value of sociality in mammalian groups. Philos Trans R Soc Lond B Biol Sci 362, 539–559. doi: 10.1098/rstb.2006.1994

Snaith, T.V., Chapman, C.A., Rothman, J.M., Wasserman, M.D., 2008. Bigger groups have fewer parasites and similar cortisol levels: a multi-group analysis in red colobus monkeys. Am J Primatol 70, 1072–1080. doi: 10.1002/ajp.20601

Vitone, N.D., Altizer, S., Nunn, C.L., 2004. Body size, diet and sociality influence the species richness of parasitic worms in anthropoid primates. Evolutionary Ecology Research 6, 183–199. doi:

Wilcove, D.S., Giam, X., Edwards, D.P., Fisher, B., Koh, L.P., 2013. Navjot’s nightmare revisited: logging, agriculture, and biodiversity in Southeast Asia. Trends Ecol Evol 28, 531–540. doi: 10.1016/j.tree.2013.04.005

Yalcindag, E., Stuart, P., Hasegawa, H., Streit, A., Dolezalova, J., Morrogh-Bernard, H., Cheyne, S.M., Nurcahyo, W., Foitova, I., 2021. Genetic characterization of nodular worm infections in Asian Apes. Sci Rep 11, 7226. doi: 10.1038/s41598-021-86518-2

